# Natural Selection Shapes the Mosaic Ancestry of the *Drosophila* Genetic Reference Panel and the *D. melanogaster* Reference Genome

**DOI:** 10.1101/014837

**Authors:** John E. Pool

**Affiliations:** Laboratory of Genetics, University of Wisconsin-Madison Madison, WI, USA, 53705

**Keywords:** Drosophila melanogaster, Drosophila Genetic Reference Panel, Admixture, Population ancestry, Linkage disequilibrium

## Abstract

North American populations of *Drosophila melanogaster* are thought to derive from both European and African source populations, but despite their importance for genetic research, patterns of admixture along their genomes are essentially undocumented. Here, I infer geographic ancestry along genomes of the *Drosophila* Genetic Reference Panel (DGRP) and the *D. melanogaster* reference genome. Overall, the proportion of African ancestry was estimated to be 20% for the DGRP and 9% for the reference genome. Based on the size of admixture tracts and the approximate timing of admixture, I estimate that the DGRP population underwent roughly 13.9 generations per year. Notably, ancestry levels varied strikingly among genomic regions, with significantly less African introgression on the X chromosome, in regions of high recombination, and at genes involved in specific processes such as circadian rhythm. An important role for natural selection during the admixture process was further supported by a genome-wide signal of ancestry disequilibrium, in that many between-chromosome pairs of loci showed a deficiency of Africa-Europe allele combinations. These results support the hypothesis that admixture between partially genetically isolated *Drosophila* populations led to natural selection against incompatible genetic variants, and that this process is ongoing. The ancestry blocks inferred here may be relevant for the performance of reference alignment in this species, and may bolster the design and interpretation of many population genetic and association mapping studies.

## INTRODUCTION

North American populations of *Drosophila melanogaster* have had a disproportionate role in classical and modern *Drosophila* genetics. They gave rise to many of the commonly used laboratory strains that came from the T. H. Morgan lab and elsewhere. More recently, the *Drosophila* Genetic Reference Panel (DGRP) [1,2] introduced a set of 205 sequenced genomes from independent inbred lines collected from Raleigh, North Carolina, USA. The DGRP has become a widely used resource for analyses of genomic variation and its relation to phenotype. Understanding the demographic history of the DGRP and other North American populations is important for maximizing the scientific value of these genetic resources. However, *D. melanogaster* is not native to the western hemisphere, and the recently arrived New World populations of this species appear to have complex origins.

*D. melanogaster* originated from sub-Saharan Africa [3], and probably from southern-central Africa in particular [4], where at some unknown time it became associated with human settlement. The species began to expand its geographic range, initially occupying more diverse environments within sub-Saharan Africa. On the order of 10,000 years ago, *D. melanogaster* managed to cross the Saharan region and expand into northern Africa and Eurasia [3,5,6]. That expansion entailed a significant loss of genetic diversity, perhaps as a result of founder event population bottlenecks, with the consequence that non-sub-Saharan populations hold only a subset of the variation observed within sub-Saharan Africa [4,5].

The expansion of *D. melanogaster* from a tropical ancestral range into temperate Old World regions appears to have had important consequences regarding adaptation to novel environments and the restriction of migration between sub-Saharan and Palearctic populations. Tropical and temperate populations have a range of morphological differences [7], and the genomic search for loci that may encode adaptive differences between these populations has been a topic of significant interest [4,8]. Partial sexual isolation has also been reported between African and non-African strains: female flies from African strains were found to discriminate strongly against non-African males [9,10,11].

North American populations of *D. melanogaster* are thought to derive from both European and African source populations. Evidence for this dual ancestry comes from three population genetic observations from North American vs. European populations. First, North American populations are more genetically similar to sub-Saharan populations than Eurasian populations are [5,12–14]. Second, these same four studies all indicate that North American populations have higher genetic diversity than European populations, an observation that seems incompatible with a simple European origin for North American populations. And third, a North American population was found to have elevated linkage disequilibrium relative to a European population [13], which is likewise consistent with recent admixture in North America. The apparent dual ancestry of North American *D. melanogaster* may have resulted from a history in which European populations initially colonized the northeast U.S., while African populations first reached the Caribbean [15,16], with subsequent geographic expansion and interbreeding leading to admixed populations.

Specific evidence that the North Carolina DGRP population is admixed comes from the analysis of Duchen *et al*. [17], who estimated these genomes to contain 15% African ancestry. That study focused on the inference of historical parameters in *D. melanogaster*, whereas the present study began by analyzing population ancestry along DGRP genomes. That analysis ultimately led to evidence that natural selection pervasively influenced the admixture process in this North American population, and continues to act on combinations of European and African alleles today. In addition to the DGRP, I also estimated ancestry along the *D. melanogaster* reference genome. Knowledge of DGRP ancestry will be important for interpreting genetic and phenotypic variation in that population, whereas mapping the European and African segments of the reference genome has important implications for the performance of reference alignment for sequenced *D. melanogaster* genomes.

## RESULTS

### Genome-wide ancestry proportions for DGRP and the timing of admixture

Non-African vs. sub-Saharan ancestry along North American *D. melanogaster* genomes was assessed using the method of Pool *et al*. [4]. This Hidden Markov Model (HMM) approach operates in genomic windows, comparing a focal genome to a non-African reference panel of genomes, and testing whether these genetic distances fit expectations based on comparisons among non-African genomes or instead match the larger distances obtained from African vs. non-African comparisons. Among the 205 DGRP genomes, inferred African ancestry averaged 19.8% with a standard deviation of 5.5% (Table S1).

From inversion-free chromosome arms, 24,034 African ancestry tracts of at least 0.05 cM were free of large-scale missing data. Based on recombination rate estimates of Comeron *et al*. [18], these tracts had a median length of 0.173 cM. Using an admixture tract length simulation approach [19] with appropriate ancestral population proportions, I estimate that this median tract size would be expected after ∼1,513 generations of admixture. The accuracy of this estimate may be affected by demographic details, natural selection, and imprecision in recombination estimates.

### DGRP ancestry proportions are highly variable along the genome

Examining collective DGRP ancestry for each window, striking genome-wide variability was detected (Figure 1). Surprisingly, the X chromosome is nearly fixed for European ancestry, with localized exceptions. Whereas the autosomes carry 21.8% African ancestry, for the X chromosome this average is reduced to 5%. 75.1% of X-linked windows have <5% African ancestry and 37.8% are completely fixed for European ancestry. In contrast, an outlier for higher African ancestry is chromosome arm 2L (Figure 1). This arm effect is largely explained by the prevalence of inversion *In(2L)t* (Figure S1), the most common African-origin inversion in the DGRP [2,20]. *In(2L)t* and other inversions can have strong effects on genetic variation across whole chromosome arms (Figure 2) [4,20].

**Figure 1.**
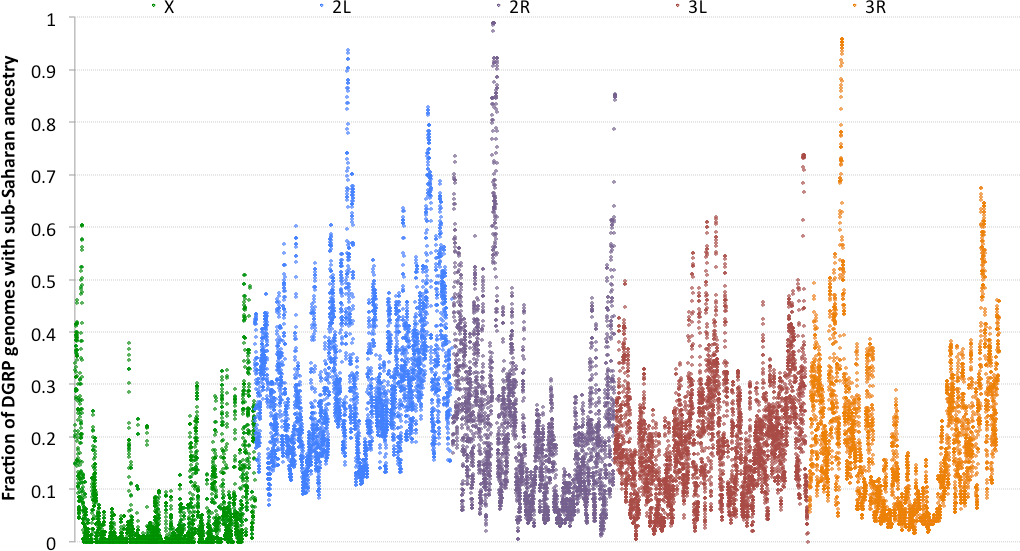
The proportion of DGRP genomes that have >50% probability of sub-Saharan ancestry in each genomic window is shown for the five major euchromatic chromosome arms (color-coded and labeled above). Genomes lacking at least 500 bp of called sequence within a given window were excluded, and windows with fewer than 50 genomes meeting that criterion were omitted from this plot.

**Figure 2.**
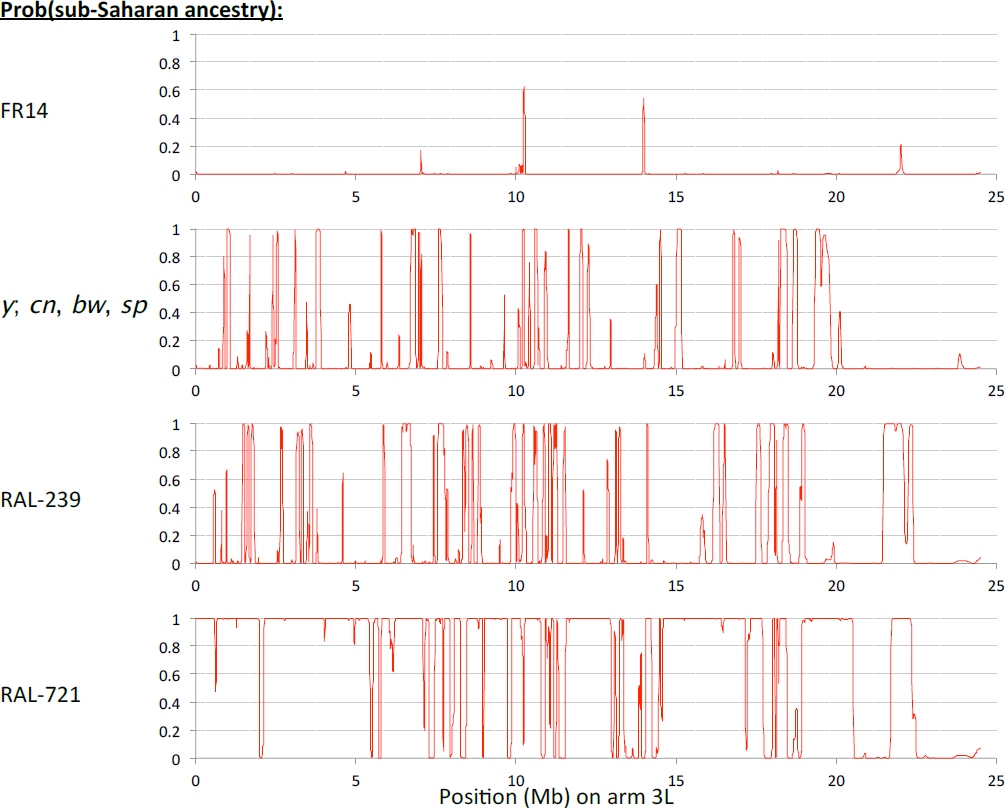
Sample ancestry likelihood plots from chromosome arm 3L show that the proportion of sub-Saharan ancestry depends on geography and inversion status. FR14 is part of the non-African reference panel, and shows almost no putative African ancestry, *y*; *cn, bw, sp* is the *D. melanogaster* reference genome; it shows peaks of African ancestry probability, often on the order of 100 kb. The DGRP genome RAL-239 shows a greater density of African ancestry tracts, but of generally similar length as the reference. Unlike the above genomes, RAL-721 carries *In(3L)P*, an inversion of sub-Saharan origin; this arm shows predominantly African ancestry with mostly narrow intervals of non-African origin.

More perplexing than the between-arm ancestry differences are the strong fluctuations observed within chromosome arms, often on the scale of tens or hundreds of kilobases (Figure 1). For each chromosome arm, there is a significant negative correlation (*P* < 0.0001) between African ancestry and recombination rate, with Pearson *r^2^* of 0.080 for standard autosomal arms analyzed jointly and 0.101 for the X chromosome. The mean sub-Saharan ancestry proportion is 30.2% for autosomal windows below 0.5 cM / Mb, but only 13.0% when the recombination rate is above 4 cM / Mb (Figure 3). This relationship is not expected under a neutral introgression scenario, but might result either from inefficient selection in low recombination regions against African alleles that are disadvantageous in the predominantly European gene pool and North American environment of the DGRP population, or from favored African alleles carrying longer linkage blocks in regions of low recombination.

**Figure 3.**
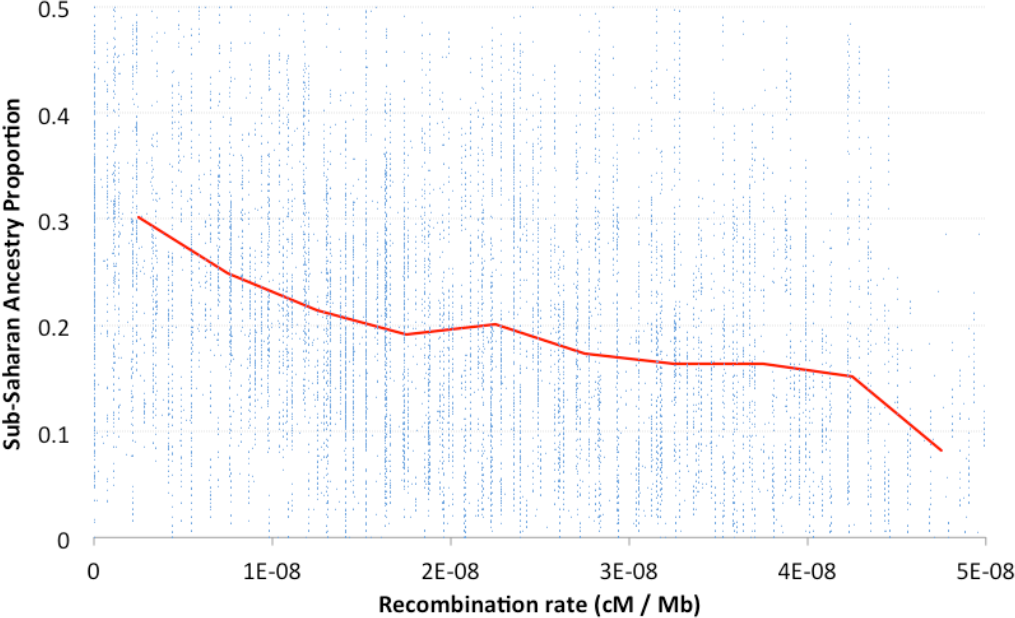
DGRP genomes display much less sub-Saharan ancestry in windows with higher recombination rates. Here, each blue dot represents one autosomal genomic window, while the red line indicates the mean sub-Saharan ancestry proportion for bins of 0.5 cM/Mb. Ancestry proportions are from standard chromosome arms only. All autosomal windows with data from at least 50 standard arms were included in statistical analyses. Only windows with recombination rate up to 5 centiMorgans per Megabase (cM/Mb) and sub-Saharan ancestry proportion up to 50% are shown in this plot.

Using simulations based on previously inferred demographic parameters, Pool *et al*. [4] found that admixture on the order of 1,000 generations ago between African and European populations of *D. melanogaster* was detected reliably by the method implemented here. The few errors tended to involve missing unusually short admixture tracts, rather than inferring false tracts. Those simulations focused on the autosomes because X-linked admixture will be easier to detect (based on the much larger diversity difference between African and European populations for the X chromosome). Hence, the lower X-linked admixture levels described above contradict the predictions of methodological bias. Nor is the recombination result easily explainable by such bias: low recombination window lengths were scaled to contain similar numbers of polymorphisms as high recombination regions, and Europe/Africa diversity ratio is typically similar across high and low recombination regions of chromosome arms [4], so there is no obvious prediction of a difference in the power to detect admixture between these categories.

### Impact of natural selection on ancestry inferences

There are reasons to be skeptical of some extreme DGRP ancestry deviations. The two intervals of maximal African ancestry are near *Cyp6g1* and overlapping *Ace*, loci with strong selective sweeps related to 20th century insecticide usage [21,22]. At these loci, sweeps that occurred after the divergence of the Raleigh population from its European and African source populations (perhaps less than 150 years ago; Keller 2007) could result in biased ancestry inference.

Although it would be desirable to annotate each case in which very recent selective sweeps may have influenced ancestry calling, this goal may require significant methodological advances. The HMM used here should be more sensitive to cases involving very recent selection affecting the European reference panel, but such sweeps could either be global (with either the same or different haplotypes fixing in each population), or shared by the European and African reference panels but not the DGRP, or specific to the European sample. These scenarios each lead to distinct predictions for variation among populations, whereas analyses focused only on the European reference panel will mainly pick up sweeps that happened prior to American colonization (which are of less concern here).

This issue reflects a general challenge for ancestry inference. Other reference panel approaches should be subject to similar effects of recent selection. Methods that do not use reference panels may return non-geographic divisions in the data, such as clustering inverted versus standard chromosome arms [20], and even if inverted arms were removed, their output could prove similarly uninformative or biased in cases of recent sweeps. Hence, the ancestry inferences presented here (Table S3) should be regarded as provisional, and should be revisited in light of future methodological developments.

For either hard sweeps or moderately soft sweeps affecting the European sample, such recent selection should increase that population’s haplotype homozygosity, since there has been very little time for mutation and recombination since the adaptive event. While such a pattern is observed at *Cyp6g1*, in general the inferred peaks of African ancestry in the DGRP show no such pattern (Figure S2). Thus, while recent selection may drive some apparent ancestry deviations, most of the genomic variance in DGRP ancestry suggested in Figure 1 may be genuine.

### Admixture in the reference genome

Using the same methods as described for the DGRP genomes, the *D. melanogaster* reference genome was estimated to have 9.4% African ancestry (Figure 2, Table S4). The relatively lesser degree of admixture in this genome, relative to the North Carolina DGRP population, would make sense if: (1) this species’ initial colonization of the New World involved an African founder population in the Caribbean and a European founder population in the northeastern U.S. [15,16], and (2) the reference strain descends primarily from laboratory stocks obtained by the T. H. Morgan laboratory in New York (as seems likely - J. A. Kennison, personal communication).

The reference genome’s segments of African ancestry are correlated with those found in the DGRP (Table 1). And like the DGRP, the reference genome is more likely to carry African ancestry in low recombination regions (Table 1). Hence, many of the demographic and selective events that molded complex patterns of ancestry in the DGRP may have affected other North American populations as well.

**Table 1.**
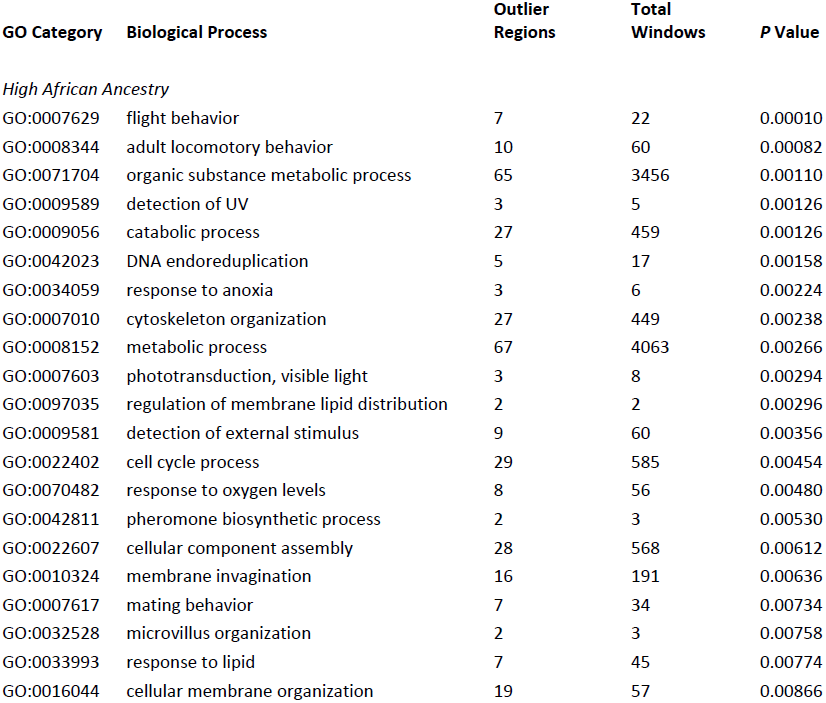
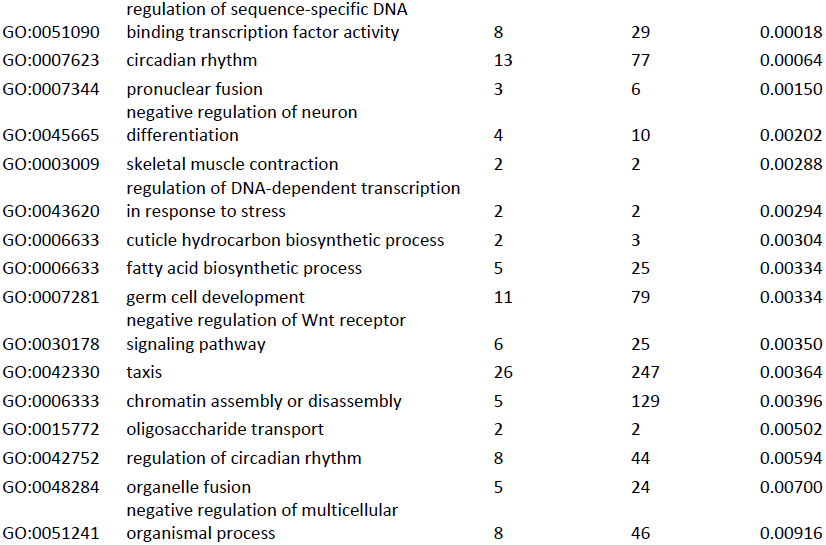
**Biological process categories over-represented in genomic outlier regions for elevated African or European ancestry**. Unique biological process GO categories with raw permutation *P* value less than 0.01 and representation in at least two separate outlier regions are shown here (with full results in Table S2). Some of these categories could reflect targets of selection favoring African or European alleles in North America, but at present they represent hypotheses for further population genetic and functional analysis.

### Functional and population genetic correlates of ancestry deviations

Although precise neutral expectations for interlocus variance in ancestry proportion depend on unknown details of the North American colonization scenario, the dramatic and non-random variance observed here suggests the possibility that African and European alleles at some loci may have had unequal fitness in North American environments. To investigate which types of genes would be the most likely targets of any such selection, gene ontology (GO) enrichment analysis was performed for intervals of elevated African or European ancestry. The GO categories most enriched for European ancestry included “circadian behavior”, while those for African ancestry included “flight behavior” and vision-related categories (Table 2; Table S2).

**Table 2.**
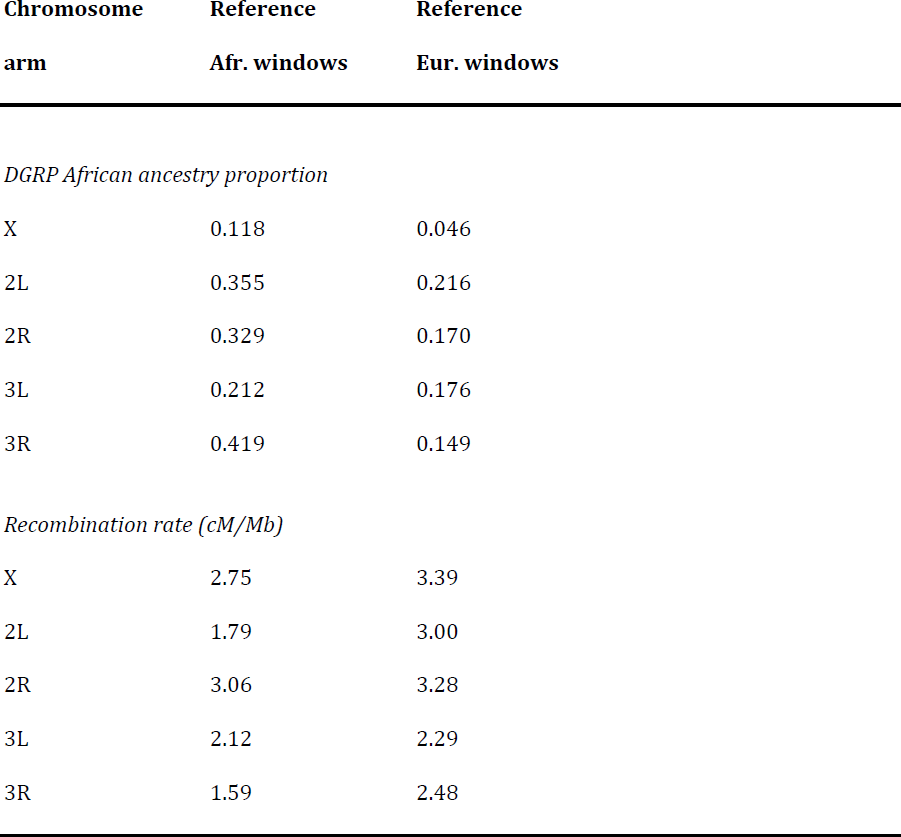
**Relationship between reference genome ancestry and DGRP ancestry or recombination rate**. For each chromosome arm, windows called as either African or European in the reference genome are compared in two respects. First, it is shown that windows called as African in the reference genome have much higher levels of African ancestry in the DGRP. Second, as observed for the DGRP, the reference genome’s African windows have lower recombination rates (based on the estimates of Comeron *et al*. 2012).

| **Chromosome arm** | **Reference Afr. windows** | **Reference Eur. windows** |
| --- | --- | --- |
| *DGRP African ancestry proportion* |  |  |
| X | 0.118 | 0.046 |
| 2L | 0.355 | 0.216 |
| 2R | 0.329 | 0.170 |
| 3L | 0.212 | 0.176 |
| 3R | 0.419 | 0.149 |
| *Recombination rate (cM/Mb)* |  |  |
| X | 2.75 | 3.39 |
| 2L | 1.79 | 3.00 |
| 2R | 3.06 | 3.28 |
| 3L | 2.12 | 2.29 |
| 3R | 1.59 | 2.48 |

### Evidence for widespread epistatic fitness interactions in the DGRP

The hypothesis that selection may disfavor certain African alleles in the primarily European gene pool of the DGRP population is consistent with the above-described relationship between recombination rate and ancestry (Figure 3). In light of previous evidence [23], at least some of these loci may be involved in interlocus fitness interactions (IFIs), in which having African alleles at one locus and European alleles at another may lead to reduced survival or fecundity. If natural selection against introgressing African alleles is ongoing today, a signal of linkage disequilibrium might be observed between pairs of loci responsible for such IFIs. Such analyses have been performed in hybridizing taxa to identify pairs of loci that may constitute Bateson-Dobzhansky-Muller incompatibilities of potential relevance to speciation [24–27].

Here, I test for disequilibrium between loci on different chromosomes, not by using individual SNP genotypes, but instead based on the ancestry calls made for each genome in each of the 24,417 genomic windows. This focus on “ancestry disequilibrium” (AD) makes genome-scale pairwise testing computationally plausible. For each interchromosomal pair of windows, I calculated a Fisher’s Exact Test (FET) *P* value, reflecting the preferential occurrence of Africa-Africa and Europe-Europe ancestry combinations at the two windows Only homozygous intervals were analyzed, so each genome has just one allele per locus, and inverted chromosome arms were excluded. Results from the true data were then compared against randomly permuted data sets, in which individual labels for the second window were shifted (thus maintaining the true data’s population ancestry frequencies at each window, as well as patterns of linkage between neighboring windows). Across the genome, a notable excess of interchromosomal window pairs with low FET *P* values was observed (Figure 4), indicating a genome-wide signal of ancestry disequilibrium. At very low *P* values, the enrichment was more pronounced for X-autosome window pairs than for pairs split between the two major autosomes (Figure 4).

**Figure 4.**
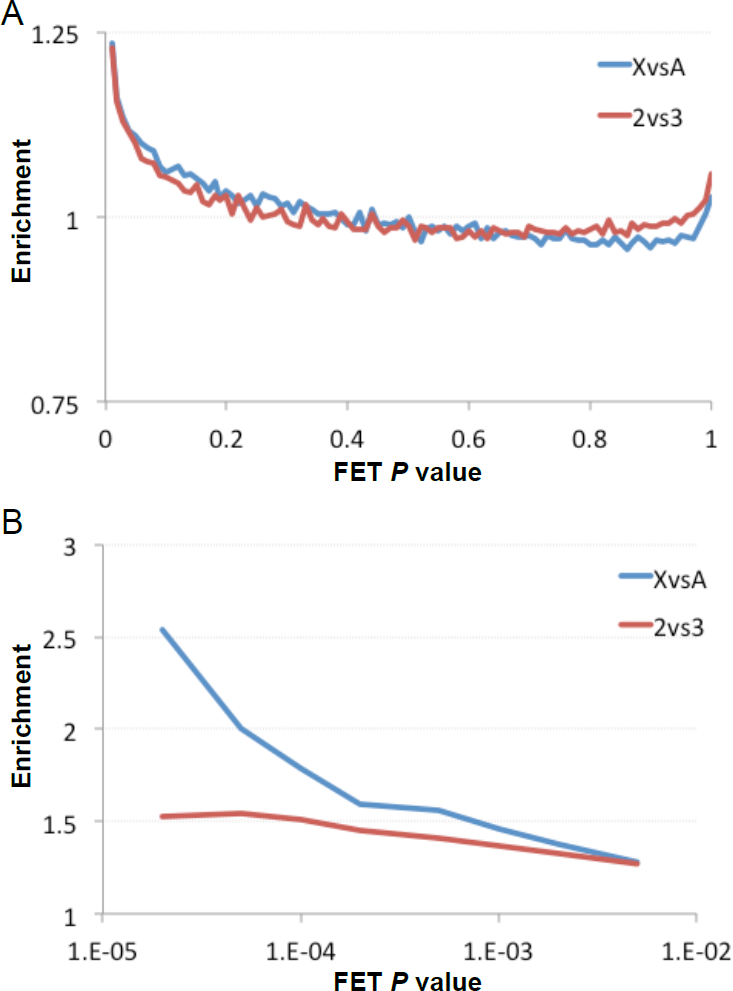
A genome-wide signal of interchromosomal ancestry disequilibrium is depicted. Here, fold-enrichment in the real data (relative to permuted data sets) is plotted for Fisher Exact Test *P* values. All comparisons between X-linked and autosomal windows, and between chromosomes 2 and 3, are plotted in separate series. Above, enrichment is plotted for each 0.01-wide *P* value bin. Below, the cumulative enrichment for all *P* values below a given threshold is indicated.

##### Box 1. Key to abbreviations used in the present study

AD: Ancestry disequilibrium – the correlation of population ancestry between loci. Analogous to linkage disequilibrium, but calculated using inferred ancestries rather than specific genotypes.
AD cluster: A pair of genomic regions that contain one or more window pairs with strong ancestry disequilibrium.
AD hub: A set of neighboring windows that overlaps an unusually large number of AD clusters. These genomic regions are hotspots for ancestry disequilibrium, and may experience interlocus fitness interactions with a number of unlinked loci.
BDMI: Bateson-Dobzhansky-Muller incompatibilities. Fitness may be compromised when variants from previously isolated populations are brought into contact by admixture.
IFI: Interlocus fitness interaction. AD may indicate an IFI, and a BDMI is a potential explanation for an IFI.
DGRP: *Drosophila* Genetic Reference Panel
DPGP: *Drosophila* Population Genomics Project
DSPR: *Drosophila* Synthetic Population Resource

To avoid treating neighboring window pairs as independent, nearby outlier *P* values were merged into two-dimensional “clusters” of ancestry disequilibrium, and these clusters were extended from each focal window until pairs with *P* < 0.05 were no longer observed with appreciable frequency. Although the binning criteria were necessarily somewhat arbitrary (see Methods), they were designed to extend clusters generously in an attempt to fully account for their effect on the genomic distribution of FET *P* values. Examining the chromosomal distribution of these pairwise clusters, there is little evidence that adjacent clusters are failing to be appropriately merged (Figure S3). This procedure resulted in 676 AD clusters with no pairwise overlap, many of which are likely to represent false positives. However, subtracting the entire span of all 676 clusters only accounted for 33% of the genome-wide excess of X-autosome FET *P* values below 0.05, and 58% of the autosome-autosome excess. Hence, although further study is needed to accurately estimate the number of pairwise IFIs between African and European alleles in the DGRP genomes, based on the present analysis I can not rule out a scenario in which a surprisingly large number of pairwise incompatibilities are present.

### Potential genetic targets of interlocus fitness interactions

The vast number of pairwise comparisons involved in genome-wide disequilibrium testing entails a multiple testing problem, with the consequence that no pairwise *P* value from a single hybrid or admixed population is likely to be statistically significant in a genome-wide context [27]. Hence, in order to draw any specific conclusions about genes causing AD in a single population, additional evidence is needed. With the goal of identifying a more confident set of AD clusters, I hypothesized that some true positive loci might participate in a greater number of pairwise interactions than expected by chance. While a plurality of all genomic windows overlapped zero AD clusters and most windows others overlapped three or fewer, a smaller subset of windows overlapped several – up to a maximum of 13 pairwise between-chromosome clusters. Comparing the total “cluster counts” of windows in the real data against those from permuted data sets, I confirmed that windows overlapping multiple pairwise clusters were observed much more frequently than expected randomly (Figure S4). For example, windows overlapping 7 or more AD clusters were 3.7X more common in the real data (implying a posterior probability of 79% that at least some of a window’s pairwise clusters are genuine), and “cluster counts” of at least 7 were observed in 59 distinct genomic regions. Windows overlapping 11 or more AD clusters were enriched by a factor of 5.2X, indicating a posterior probability of 84%. Hence, a subset of windows constituting “AD hubs” have fairly strong confidence of holding genuine IFIs, even though data are limited to just one admixed population.

AD in North American *D. melanogaster* could indicate IFIs resulting from adaptive functional differences between the African and European source populations. For a gene where natural selection had acted differently between those two populations, we might expect locally elevated *F_ST_* values between European and West African populations. Consistent with this hypothesis, windows overlapping the largest number of AD clusters were somewhat more likely to have high *F_ST_* values between European and West African populations (Figure S5). Importantly, *F_ST_* peaks are typically narrower than AD hubs, so their co-occurrence may help to localize the genetic targets of IFIs.

A thorough analysis of genes likely to underlie IFIs in North American *D. melanogaster* could encompass one or more follow-up studies. Still, a preliminary examination of the genes and pairwise combinations involved in AD hubs may motivate hypotheses for further genomic and functional testing, regarding the biological nature of putative incompatibilities between African and European *D. melanogaster*. I therefore highlight a few of the most notable genes and categories indicated by these AD hubs below.

Figure 5 illustrates the pairwise components of AD hubs with at least 7 pairwise interchromosomal interactions. The most extreme AD hub, overlapping 13 clusters, was centered on the gene *Argonaute 2* (Figure 6). An RNA interference gene, *AGO2* is involved in the loading of siRNA onto the RISC complex, and its known functions include antiviral response, chromatin silencing, and autophagy. Along with a second AD hub including *Dicer-2*, this result is consistent with the previous finding that RNAi genes are frequent targets of positive selection in *Drosophila* [29]. Another previously implicated target of selection is *polyhomeotic-proximal* (*ph-p*), which sits within one of just two AD hubs that overlap 12 clusters. This polycomb group gene, which has roles in gene silencing, nervous system development, and ecdysone response, was previously shown to have experienced a selective sweep in the African ancestral range [30], and may have experienced additional adaptation outside Africa as well.

**Figure 5.**
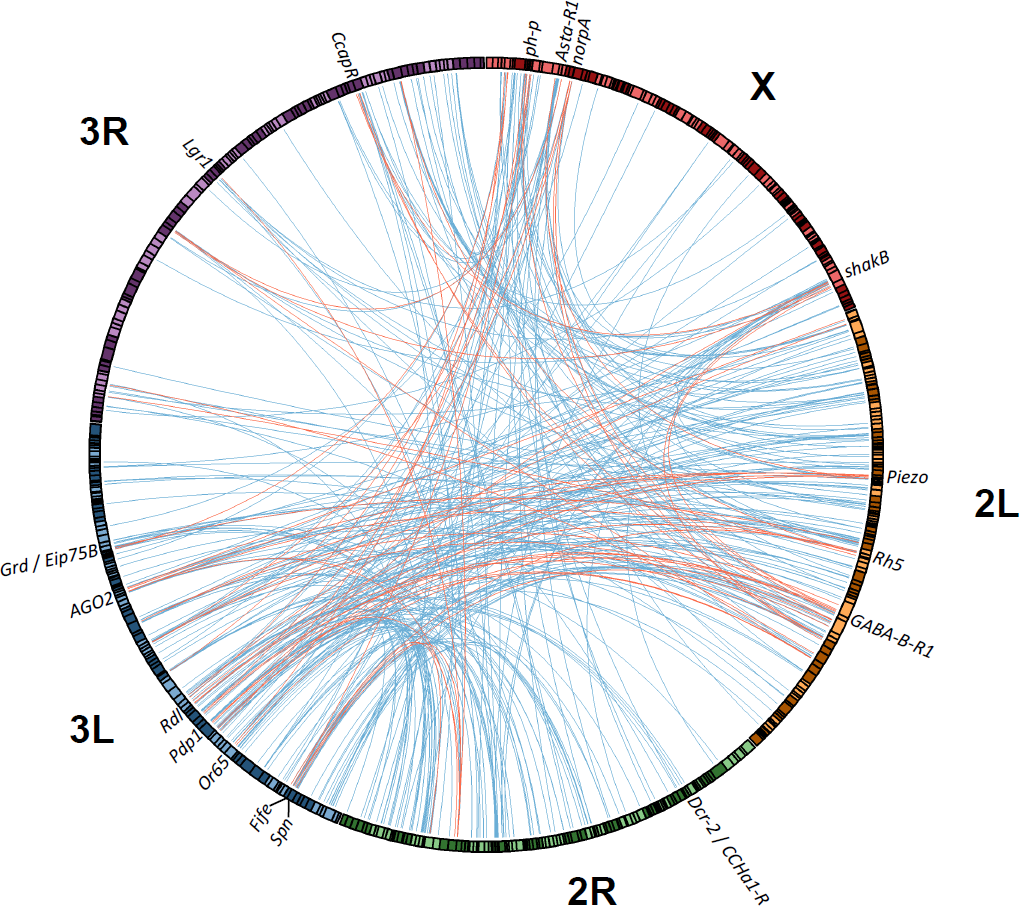
Interactions involving ancestry disequilibrium hubs that contain elevated Africa-Europe *F_ST_* (see Methods] were plotted using Circos [28]. AD clusters linking two unlinked AD hubs are shown in red, while those involving a single AD hub are in blue. Locations of genes mentioned in the text are shown; these genes were indicated by patterns of cluster overlap and *F_ST_* but further research is needed to assess their potential involvement in interlocus fitness interactions.

**Figure 6.**
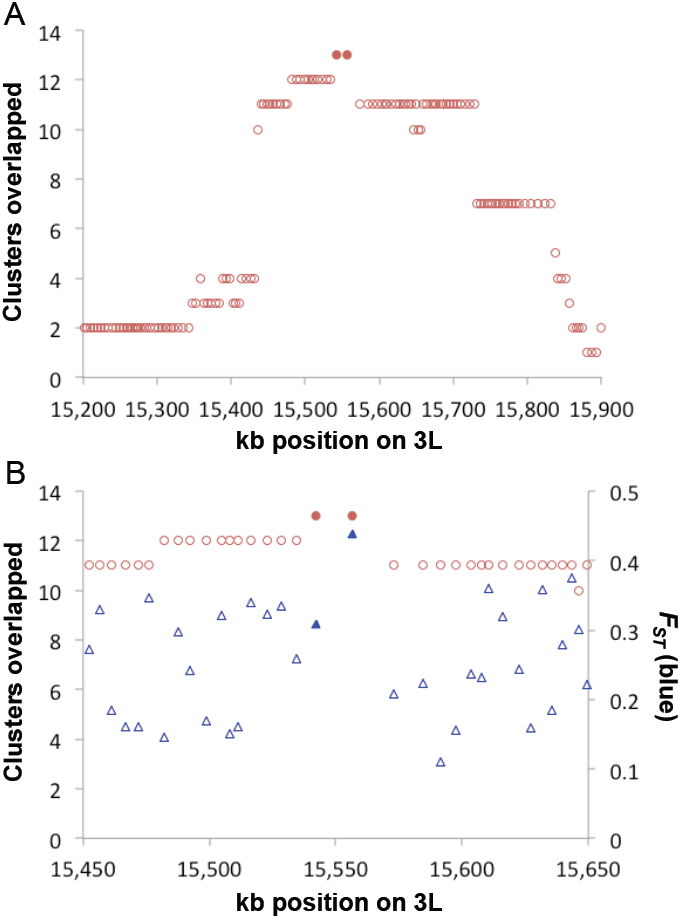
The co-occurrence of a strong ancestry disequilibrium hub and elevated *F_ST_* at the gene *Argonaute 2*. (A) The two windows overlapping *AGO2* are shaded; these represent the largest number of pairwise AD clusters overlapping any windows in the genome. (B) The same cluster overlap statistic is plotted across a narrower region, alongside window *F_ST_* between European and western African populations. High values of *F_ST_* could indicate adaptive functional differences between the two source populations of the North American DGRP population (the median *F_ST_* on 3L is 0.19). Nearly all of the *AGO2* transcript is located within the right-hand shaded window with elevated *F_ST_*.

AD hubs displaying the greatest number of interactions (≥11) and elevated *F_ST_* values also included the genes *Allatostatin A receptor 1* (*AstA-R1*; neuropeptide signaling), *Fife* (a recently-described regulator of synaptic transmission [31]), and *shaking B* (*shakB*; a synaptic gap junction protein involved in phototransduction). Other genes in AD hubs (≥7 clusters) with elevated Africa-Europe *F_ST_* values included additional neuropeptide and hormone receptors (*e.g*. the unlinked GABA receptors *Rdl*, *GABA-B-R1*, and *Grd*, along with *Eip75B*, *CCAP-R*, *CCHa1-R*, and *Lgr1*), plus other genes involved in phototransduction and/or circadian rhythm (*norpA*, *Pdp1*, *Rh5*), as well as other genes involved in olfaction and sensory behavior (*Piezo*, *Spn*, the *Or65* cluster). A full description of AD hubs and their associated interactions is given in Table S5.

Enriched GO categories for windows in AD hubs with elevated *F_ST_* (see Methods) echoed many of these same themes (Table S6). These categories included “detection of chemical stimulus involved in sensory perception” (which had the lowest *P* value among GO categories represented by at least 5 AD hubs), “cellular response to stimulus”, “signal transducer activity”, “cell surface receptor signaling pathway”, “intrinsic to membrane”, aspects of transmembrane transport, and GABA and allatostatin receptor activities. Windows from 35 AD hubs met the *F_ST_* criteria for this analysis, encompassing a median span of just 10 kb per hub. For 27 of these hubs, the window(s) with elevated *F_ST_* included at least one gene from the GO categories mentioned above. This exploratory analysis can not conclusively point to the genes and processes underlying putative incompatibilities in the DGRP, but it does suggest hypotheses for downstream molecular and genomic studies.

Less than a third of pairwise clusters involving AD hubs linked one hub to another (Figure 5). While some of these two-hub interactions could make sense based on related known functions (*e.g*. a cluster that links *shaking B* and *Pdp1*), other interactions are less functionally obvious (*e.g. AGO2* with *Piezo*, and with *Rh5*). The fitness interactions implied by AD need not involve direct molecular interactions; they could instead stem from higher-order phenotypes. Still, AD analysis may be fairly unique among evolutionary genomic methods in its potential to identify novel functional relationships between genes. This signal could complement other genomic searches for interactors, such as correlated rates of protein evolution among taxa [32]. But notably, AD can operate on a shorter time scale, it only requires data from a single species, and is not confined to interactions between protein-coding sequences.

## DISCUSSION

### Evolutionary history and genetic composition of North American D. melanogaster

North American strains of *D. melanogaster*, including the reference genome strain and the DGRP, have taken on great importance in a wide range of genetic studies. Previous studies had suggested that New World populations are likely to have an admixed history, descending from source populations both in Europe and in the sub-Saharan ancestral range [12,15]. However, this complex history and dual ancestry, along with its significance for *Drosophila* research, have not been broadly appreciated in the literature, and efforts to study the genetic composition of important fly strains have been lacking.

In this study, I estimated population ancestry (European or African) along the genomes of the reference strain and the DGRP. Results strongly support the hypothesis of admixture between these source populations in North America. The larger minority of African ancestry in the mid-Atlantic DGRP (20%) versus the northeastern reference genome (9%) is consistent with the possibility that an ancestry gradient exists among U.S. populations, with greater European ancestry in the north and somewhat higher proportions of African ancestry in the south, perhaps as a consequence of secondary contact after two separate colonizations.

### *An empirical estimate for the generation time of* D. melanogaster

By combining our estimate of the timing of admixture (1,513 generations, based on the recombination-mediated shortening of ancestry tracts over time) with historical records on the American expansion of *D. melanogaster*, I can obtain a rough empirical estimate of the average generation time of this population. In light of the hypothesized colonization of the New World by European strains via the northeast U.S. and by African strains via the Caribbean [15,16], one plausible site of initial secondary contact would be in the far southeastern U.S. Based on the first observation of *D. melanogaster* from the southern U.S. (1894 in Florida [16]), 109 years would have elapsed before the collection and inbreeding of the DGRP strains in 2003. Thus, an estimate of the number of generations per year is 1,513 / 109 = 13.9. Our estimate of 1,513 generations since admixture carries important uncertainties (see Results), as does the precise timing of secondary contact. Furthermore, generation time may vary geographically and temporally based on climate and other factors. Temperate populations are known to undergo reproductive diapause in winter [33,34]; Raleigh’s climate would seem unfavorable for outdoor reproduction of *D. melanogaster* for roughly three months of the year, whereas other populations may have longer or shorter reproductive seasons based on temperature, rainfall, resource availability, and other factors. In spite of these caveats, the above estimate may be preferable to the commonly used figure of 10 generations per year, for which no empirical basis is typically cited. Improving our estimates of this quantity is important, because it represents a critical parameter for relating DNA variation and evolution to historical time, whether one is studying changes on the scale of months or millions of years.

### Genome-wide evidence for natural selection shaping patterns of admixture

Three primary patterns in the DGRP ancestry inferences suggest that natural selection has powerfully influenced patterns of population ancestry along these genomes. First, levels of European and African ancestry vary strikingly within and between chromosome arms (Figure 1). Second, the degree of African introgression is greatly reduced in regions with higher recombination rates (Figure 3). Third, there is a genome-wide abundance of interchromosomal AD locus pairs in which strain ancestries are correlated (Figure 4).

With regard to the first point, one striking feature of the genomic ancestry landscape is the X chromosome’s strongly reduced African introgression relative to the autosomes. This result mirrors the situation in sub-Saharan Africa, where admixture from outside Africa is lowest on the X [4,35]. X chromosomes may thus be inhibited from introgressing between African and non-African populations in either direction. Qualitatively similar patterns have been reported from cases of hybridization involving mice, Neanderthals, and other taxa [36–38]. Although the present results concern the admixture of two populations of the same species, they are compatible with Haldane’s Rule [39]. They are also concordant with the findings of Lachance and True [23], who reported a substantial rate of epistatic fitness interactions between X-linked and autosomal loci in crosses involving Canadian and Caribbean strains. Haldane’s Rule suggests an elevated contribution of the X chromosome to reproductive isolation, potentially involving Bateson-Dobzhansky-Muller incompatibilities (BDMIs) in which between-locus combinations of alleles that had never co-existed arose following population divergence. A greater effect of BDMIs involving X-linked loci could be explained by recessive BDMIs (which are readily exposed in hemizygous males), or by a greater density of functional differences between African and European populations on the X chromosome, among other potential explanations [40].

Patterns of ancestry within chromosome arms further suggest that European and African alleles at many loci have had unequal fitness in the DGRP population. Sharp peaks of African or European ancestry were apparent. Levels of African introgression into this primarily European population were strongly dependent on recombination rate, with much less African introgression in higher recombination regions. Part of this signal could result from African alleles favored in the DGRP population, with longer linkage blocks hitchhiking with them in regions of low recombination. Alternatively, in accord with the X-autosome contrast and the ancestry disequilibrium analysis, African alleles at many loci may have been selected against in the DGRP population (perhaps due to incompatibilities with European alleles at other loci, or else directional selection based on the North Carolina environment or the prevalent mating system). In either case, the functional differences between African and European populations acted upon by selection in North America are likely to have been products of positive selection in one or both of the source populations. The brief evolutionary time scale of these populations’ separation (perhaps only ∼0.06*N*_*e*_ generations [6]) leaves little time for mutation and drift alone to produce such differences.

### Ancestry disequilibrium and its possible causes

Consistent with the hypothesis of epistatic incompatibilities or other fitness interactions between African and European alleles, I found that ancestry disequilibrium is widespread in the DGRP genomes and may involve a large number of locus pairs. The most obvious explanation for AD is an incompatibility between an African allele at one locus and a European allele at another, producing an epistatic fitness interaction due to consequences for survival and/or reproductive success. Positive assortative mating - if flies with African alleles at certain loci mate preferentially – might also contribute to ancestry disequilibrium among wild-caught individuals. Thus, AD could stem from interactions between individuals in addition to epistasis within individuals. It is worth mentioning, however, that the present study does not directly examine wild-caught flies, but instead the genomes of strains that were inbred for 20 generations, and had originated from >200 independent isofemale lines. Recessive BDMIs will be unmasked by the inbreeding process. Although opportunities for natural selection are limited during inbreeding, the success of a full sibling cross might be influenced by the combinations of African and European alleles that these individuals possess. Thus, inbreeding and the opportunity to study mostly-homozygous genomes may amplify the signal of IFIs and aid the search for causative loci.

Another recent study that used an inbred *Drosophila* collection to test for genetic incompatibilities within *D. melanogaster* was by Corbett-Detig *et al*. [41]. These authors used genotyping data from the Drosophila Synthetic Population Resource (DSPR) [42], which consists of more than 1,700 recombinant inbred lines from panels that derive from 8 geographically diverse founder strains after 50 generations of interbreeding. Corbett-Detig *et al*. [41] found evidence of interchromosomal allelic associations, concluding that they stemmed from incompatibilities segregating within populations. In light of the current study, and given the mix of cosmopolitan and sub-Saharan strains in the DSPR, it is also possible that some of these incompatibilities had accumulated between populations. Examining the 22 SNP pairs identified by that study, none of the corresponding window pairs had evidence for AD in the present analysis (*P* > 0.05 in all cases where African alleles existed at both loci), and only 1 of the 44 SNPs was located within an AD hub (a window on arm 3R with Europe - West Africa *F_ST_* of 0.46, upstream of the genes *βTub97EF* and *CG4815*). However, the 30 autosomal windows containing these SNPs had a median ancestry deviation statistic (see Methods) of 5.8% toward European ancestry (Mann-Whitney *P* = 0.0037 comparing these SNPs’ ancestry deviations against all other autosomal windows). This pattern could indicate that at least some of the SNP’s identified by Corbett-Detig *et al*. [41] might be unrelated to Africa-Europe genetic differentiation. Alternatively, one could suppose that these DSPR loci were subject to strong epistatic selection (to be observed on a short laboratory time scale), and that such alleles might have been purged from the DGRP population by now, while AD in the present study may be driven by incompatibilities of more moderate effect. More generally, further analysis of the selection coefficients that may drive DGRP ancestry deviations and ancestry disequilibrium, in light of nuances such as demographic details and dominance, is warranted.

Previously, it was shown that one solution to the multiple testing problem inherent in genome-wide disequilibrium testing is to add data from a second independent hybrid / admixed population and require that both populations show a disequilibrium signal for a given pair of loci [27]. Appropriate genomic data is not yet available for such an analysis in *D. melanogaster*, since the admixture tracts found in sub-Saharan populations are still impractically long for locus-specific analysis [4]. However, the two-population approach could become feasible if a number of strain-specific genomes were sequenced from a region such as Saharan Africa, Madagascar [5], northern Australia, or possibly South America. The suitability of a population will depend on the timing and amount of admixture, and analysis supporting an independent history of admixture relative to North America.

Here, I proposed that without data from a second population, statistical power can be gained by focusing on “AD hubs”. Indeed, I found that loci participating in multiple pairwise interactions were far more common in the real data than expected for random false positives. This step allows the identification of a set of loci with fairly strong confidence of contributing to IFIs (*e.g*. 79% to 84% posterior probabilities), including those discussed above. Many of these AD hubs include genes with roles in neurotransmission and sensation. It is not possible to infer from the present data what caused fitness interactions involving this group of genes, whether it be ecological or reproductive aspects of behavior, the maintenance of function in novel thermal environments, or other selective pressures. Such hypotheses will ultimately require experimental analysis.

It will also be of interest to compare the genomic admixture patterns identified in the DGRP to broader latitude clines in eastern North America and elsewhere [43–45], with the expectation that many loci subject to ancestry deviations or ancestry disequilibrium in this North Carolina sample may show atypical clinal patterns as well. However, such analyses should ideally be conducted based on ancestry proportions along the cline, as opposed to *F_ST_* between northern and southern populations. If a latitude gradient in ancestry is present, then heterogeneity in north-south *F_ST_* may simply reflect a ragged genomic landscape of genetic differentiation between the African and European source populations. Since genomic data from *D. melanogaster* latitude clines mainly comes from pooled sequencing, a method to estimate ancestry proportions from pooled data would allow for more robust clinal analysis.

I have not estimated the precise number of loci contributing to ongoing fitness interactions in the DGRP population, and further methological advances toward this goal would be desirable. However, the above analyses hint that this number may be substantial. Excluding several hundred of the most extreme pairwise interactions did not erase the genomic signal of AD. The identification of 59 AD hubs at a 79% confidence level is relevant as well, as is the observation that these hubs appear to interact with a larger number of partner loci (Figure 5). These findings, together with the pronounced genomic variance in ancestry and its correlation with recombination rate, suggest that natural selection has profoundly altered the genomic consequences of admixture between temperate and tropical populations of *D. melanogaster*. This work provides an intriguing example of admixture between genetically differentiated populations, in a species in which large populations may facilitate an important role for natural selection in the genome [46,47]. Importantly, this may also be a system in which putative incompatibilities are particularly amenable to functional characterization.

### *Significance of mosaic ancestry for* Drosophila *research*

Being the first and most completely sequenced *D. melanogaster* genome, the genome of the *y*; *cn, bw, sp* laboratory strain is typically the standard against which newly sequenced genomes from this species are compared. In an evolutionary context, however, this genome is not an obvious “reference”, being the result of a complex history involving founder events and admixture. The reference genome’s mosaic ancestry may impact reference alignments and downstream analyses. Non-African *D. melanogaster* have essentially a subset of the genetic diversity present in sub-Saharan Africa. Thus, a pair of non-African genomes will have fewer sequence differences than a pair of sub-Saharan genomes or a comparison between these groups. During reference alignment, too many SNP or indel differences from the reference genome may cause reads not to map. Thus, when the reference carries a European allele, reads from other non-African alleles may have a higher probability of mapping than reads from sub-Saharan alleles. This effect may depend on the method and parameters used, but could bias population genomic studies of individual genomes or pooled samples in ways that are heterogeneous across the genome. This problem might be minimized by accounting for known variation during reference alignment, or by using a reference genome with similar genetic distances to all strains of *D. melanogaster* (*e.g*. from Zambia [4]).

The mosaic ancestry of DGRP and laboratory strains may also be relevant to a range of phenotypic and genetic studies. The European and African source populations probably differed in various phenotypes [7]. Some of the phenotypic diversity resulting from their admixture may persist today and contribute to the trait variation of populations such as the DGRP. As a potential example, variants at many of the AD hub genes mentioned above were found to have associations with sleep traits in the DGRP [48], including *AlstA-R1*, *βTub97EF*, *Eip75B*, *Grd*, *norpA*, the *Or65* cluster, *Rdl*, *shaking B*, and *Spn*. It could be worthwhile to incorporate ancestry into similar genome-wide association studies. Ancestry-associated phenotypic variation might have longer linkage blocks flanking the causative sites, potentially making it easier to detect (if this linkage signal could be incorporated), but possibly more challenging to localize. In light of the ancestry disequilibrium results cited above, admixture could also be a source of epistatic interactions in the DGRP and lab strains, which might impact phenotypes of interest and contribute to genetic background effects.

## MATERIALS AND METHODS

### Genomes and ancestry inference

Aside from the *D. melanogaster* reference genome (release 5.57), all genomes analyzed here were originally described in the Drosophila Genetic Reference Panel (DGRP) [1,2] or the Drosophila Population Genomics Project (DPGP), phase 2 [4]. The alignments used in the present study were generated using a common pipeline, involving a second round of mapping to a modified reference genome, as described by Lack *et al*. [46].

Ancestry estimation was performed using the Hidden Markov Model (HMM) approach originally described by Pool *et al*. [4]. Briefly, this method utilizes the difference in genetic distance between two types of pairwise comparisons: (1) comparisons among “cosmopolitan” genomes from outside sub-Saharan Africa, which have reduced diversity stemming from the out-of-Africa bottleneck, and (2) comparisons between sub-Saharan and cosmopolitan genomes, which have similarly higher distances as comparisons between sub-Saharan genomes. These comparisons are evaluated with the aid of two reference panels of genomes (sub-Saharan and cosmopolitan). Distances are initially assessed in non-overlapping windows across the genome. In each window for a focal genome being tested, its genetic distance to the cosmopolitan panel is tested to evaluate whether it more closely resembles the comparisons among cosmopolitan genomes (perhaps indicating cosmopolitan ancestry) or the comparison between the sub-Saharan and cosmopolitan genomes (favoring sub-Saharan ancestry of the tested genome). A likelihood of each ancestry type is obtained for each window for this genome, with the HMM then returning final ancestry probabilities in each case.

Following Lack *et al*. [49], the sub-Saharan reference panel consisted of 27 Rwanda genomes, while the cosmopolitan panel included 9 France and 3 Egypt genomes. Chromosome arms with inversions were excluded from reference panels, based on evidence that inversions have recently moved between populations [4,20]. Based on the relatively older admixture of North American populations (compared with the apparently very recent introgression studied in Africa), a somewhat smaller window size was used in the present analysis. Windows were scaled by genetic diversity, as defined by 100 non-singleton SNPs in the Rwanda sample. In moderate to high recombination regions, these windows typically corresponded to 3-5 kb. Otherwise, ancestry was assessed exactly as previously described [4,49]. Regions of genomes previously inferred to contain residual heterozygosity or identity by descent with another analyzed genome were excluded from all analyses.

### Ancestry deviations and gene ontology enrichment

Population ancestry proportions among DGRP genomes were found to vary on both local and broader genomic scales. To analyze genes that could be responsible for local peaks of African or European ancestry, a simple “ancestry deviation” statistic was implemented. This statistic was defined as the difference between the proportion of African ancestry in the focal window and the median of that quantity in the 51st to 250th windows on each side. This procedure helped to account for the regional ancestry background while excluding windows that may deviate along with the focal window due to the same instance of natural selection. Outlier windows for ancestry deviation were defined as based on the 2.5% tails for each chromosomal arm. To avoid double-counting the same putative instance of selection, “outlier regions” grouped outlier windows with up to two non-outlier windows between them.

The set of all genes overlapping outlier regions (including the next exon on each side of the region) was subjected to gene ontology (GO) enrichment analysis. GO categories corresponding to the overlapping genes were counted only once per region. The locations of all outlier regions (in terms of the windows that each spanned) were randomly permuted within their original chromosome arms 50,000 times, a practice that accounts for the effects of varying gene lengths. For each GO category, the proportion of random permutations generating at least as many outliers as observed in the real data constituted a *P* value.

### Ancestry disequilibrium testing and analysis

Analogous to linkage disequilibrium, I tested for “ancestry disequilibrium” (AD) using the ancestry inferred for each genome in each window, asking whether having an African allele in one window boosted the chance of having an African allele in an unlinked window. Fisher Exact Tests (FETs) were applied to each interchromosomal pair of windows. Genomic distributions of FET *P* values were compared between the real data and permuted data sets in which individual labels were consistently shifted for the second window in a pair (thus maintaining linkage patterns among windows). Due to the computationally intensive analysis, just 10 permuted data sets were assessed, but each one contains roughly 1 E +8 *P* values, and consistent results were observed from one replicate to the next.

To bin multiple neighboring window pairs that could result from the same pair of interacting loci, a set of the most extreme AD window pairs were extended to form “AD clusters”. Specific criteria for selecting and extending these criteria were as follows. (1) Identify each interchromosomal window pair with a raw FET *P* value below 0.0001 as starting points for AD clusters. (2) While holding one member of the focal window pair constant, extend the cluster from the other window by advancing in each direction until 10 consecutive *P* values above 0.05 are observed. Repeat to extend bidirectionally from the first member of the window pair as well, holding the second member of the window pair constant. (3) Consider clusters to encompass the full two-dimensional range of windows between the window start and stop positions identified for each side of the pair above. Merge any clusters that have overlapping boundaries on both chromosomes, giving the merged cluster the maximal span indicated by the boundaries of its component clusters.

GO enrichment analysis on AD hubs with elevated Africa-Europe *F_ST_* [50] was conducted as described above for African ancestry deviation, except for the specific criteria for outlier regions. Windows were considered to lie within AD hubs if they overlapped at least 7 interchromosomal AD clusters, and if their cluster count was within 1 of the local maximum (with cluster counts of 4 or below preventing the extension of AD hubs). To focus on loci with at least modest evidence for adaptive differences between the source populations of North American *D. melanogaster*, a minimum *F_ST_* of 0.35 (autosomes) or 0.42 (X) was required, corresponding to roughly the upper 15% quantile of this statistic. *F_ST_* was evaluated between a France sample and a panel of four small western African samples from Cameroon, Gabon, Guinea, and Nigeria [4].

## ACKNOWLEDGEMENTS

I thank J. J. Emerson for assistance with the admixture detection HMM, R. B. Corbett-Detig for helpful manuscript suggestions, and J. B. Lack for bioinformatic assistance.

## SUPPORTING INFORMATION CAPTIONS

**Figure S1.** The proportion of non-inverted DGRP chromosome arms that have >50% probability of sub-Saharan ancestry in each genomic window. Aside from the exclusion of inverted chromosome arms, this analysis is identical to that shown in Figure 1. By comparison, this plot shows less broad-scale elevation of sub-Saharan ancestry on arm 2L (except near the centromere), reflecting the exclusion of genomes carrying *In(2L)t* and its corresponding African ancestry block. Regions of particularly low sub-Saharan ancestry are now more apparent on arms 2R and 3R, reflecting the removal of inversions *In(2R)Ns* and *In(3R)P*, which also have sub-Saharan origin (Corbett-Detig and Hartl 2012).

**Figure S2.** Outlier windows for high African ancestry in the DGRP do not generally have an excess of identical haplotypes in the non-African reference panel. If the France/Egypt panel had experienced very recent selective sweeps (since the specie’s New World colonization), spurious inference of African ancestry would be possible. However, such recent selection would be expect to produce long identical haplotypes, which are rare across most of the *D. melanogaster* genome. Above, it can be observed that loci with higher African ancestry proportions in the DGRP tend not to have low values for the haplotype heterozygosity quantile in the non-African reference panel (the proportion of windows with lower or equal haplotype heterozygosity, calculated separately for each autosomal arm). This result suggests that most putative peaks of African ancestry are unlikely to be an artefact of recent selective sweeps.

**Figure S3.** Ancestry disequilibrium clusters are scattered across chromosome arms. Here, the midpoints of each AD cluster linking chromosome arms 2L and 3L are plotted (the pair of arms with the greatest number of clusters). If multiple AD clusters were often caused by the same underlying AD signal, points in the above plot would often be tightly grouped. Although there are isolated cases above where two or more clusters might share the same basis, the overall patterns suggests that most AD clusters - whether they represent true or false positives - appear to represent distinct signals in the data. Future statistical methodological development should target the refinement of two-dimensional AD regions.

**Figure S4.** Cumulative enrichment and statistical confidence of ancestry disequilibrium hubs are plotted. “Enrichment” (*e*) refers to the fold-excess of windows overlapping *X* or more clusters in the real data, relative to randomly permuted data sets. Posterior probability is calculated as *e* / (1 + *e*). Very similar results when windows in low recombination regions were excluded (not shown).

**Figure S5.** The average *F_ST_* for windows overlapping 10 or more AD clusters is compared to that for all other windows, with the autosomes and X chromosome analyzed separately. Here, *F_ST_* is measuring genetic differentiation between European and western African populations, as proxies for the source populations that gave rise to North American *D. melanogaster*. Elevated *F_ST_* for these hubs is consistent with the hypothesis that adaptive functional differences had arisen between the source populations, which may then have been subject to interlocus fitness interactions upon secondary contact in the New World.

**Table S1: Estimated proportion of European (cosmopolitan) ancestry in each of the 205 DGRP genomes**.

**Table S2: Gene Ontology Categories Enriched for African or European Ancestry**. The following two tabs summarize results for GO category enrichment in outlier regions of African and European ancestry, respectively. Details concerning the methodological approach are provided in the Methods section of the main paper.

**Table S3: Intervals estimated to have >50% probability of sub-Saharan ancestry from the DGRP genomes**. Start and stop positions (Flybase release 5 numbering) are presented in separate tabs for each of the five major euchromatic chromosome arms.

**Table S4: Intervals estimated to have >50% probability of sub-Saharan ancestry from the reference genome**. Start and stop positions (Flybase release 5 numbering) are presented for each of the five major euchromatic chromosome arms.

**Table S5: Genomic locations of ancestry disequilibrium hubs and their corresponding interactions**. In the following tabs, summaries are given of all AD hubs (overlapping 7 or more AD clusters), and the subset of those AD hubs with elevated *F_ST_* (see methods). Additional tabs describe all AD clusters involving at least one AD hub.

**Table S6: Gene ontology categories enriched for ancestry disequilibrium hubs with elevated genetic differentiation**. This analysis concerns windows within the cluster count peak of an ancestry disequilibrium hub, that also have elevated genetic differentiation between European and Western African populations. For each gene ontology category, the following table indicates its statistical enrichment among the genes overlapping these windows, compared to random expectations. Details concerning the methodological approach are provided in the Methods section of the main paper.

